# Haplotype block dynamics in hybrid populations

**DOI:** 10.1101/058107

**Authors:** Thijs Janzen, Arne W. Nolte, Arne Traulsen

## Abstract

When species originate through hybridization, the genomes of the ancestral species are blended together. Over time genomic blocks that originate from either one of the ancestral species accumulate in the hybrid genome through genetic recombination. Modeling the accumulation of ancestry blocks can elucidate processes and patterns of genomic admixture. However, previous models have ignored ancestry block dynamics for chromosomes that consist of a discrete, finite number of chromosomal elements. Here we present an analytical treatment of the dynamics of the mean number of blocks over time, for continuous and discrete chromosomes, in finite and infinite populations. We describe the mean number of haplotype blocks as a universal function dependent on population size, the number of genomic elements per chromosome, the number of recombination events, and the initial relative frequency of the ancestral species.

---

Speciation through hybridization has long been recognized as a potential driver in the formation of new species in plants (Grant 1981). More recently, it has also received attention as a process that may lead to speciation in animals (Abbott *et al.* 2013). It has been pointed out that genetic admixture between differentiated lineages should only be considered hybrid speciation when the joint contribution of both parental species is instrumental in the rise of the new species, for example by creating direct barriers to reproduction with the parental species or by facilitating ecological isolation of the emerging hybrid lineage (Mallet 2007; Nolte and Tautz 2010; Abbott *et al.* 2013). Hybrid speciation involves that parental genetic variance that reduces the fitness of the emerging lineage is purged or selected for if it helps to adapt to a new niche (Buerkle et al. 2000; Barton 2001). Although these studies predicted a lag phase during which a hybrid lineage has to go through an evolutionary optimization, empirical studies suggest that hybrid speciation can occur rapidly, possibly within hundreds of generations (Nolte *et al.* 2005; Buerkle and Rieseberg 2008; Abbott *et al.* 2013). Hence, systems and methods to gain better insight in the timeframes required for hybrid speciation are needed. Conventional molecular clock estimates are usually too coarse to be applied to cases of rapid speciation, but lineages of hybrid origin hold the potential to estimate rather short time-frames from the ancestry structure of admixed genomes (Buerkle and Rieseberg 2008; Liang and Nielsen 2014). Newly formed hybrids contain contiguous genomic blocks that originate from either one of the ancestral species and decay from generation to generation through genetic recombination. Understanding how ancestral genomic blocks decay over time can inform us about how genomes of hybrid lineages evolved.

Fisher already recognized that the mix of genetic material after a hybridization event is organized within contiguous haplotype blocks. The dynamics of the delineation between these blocks, ‘junctions’, can be traced through time, and he formulated the expected number of junctions given the number of generations passed since the onset of hybridization (Fisher 1949, 1954). Fisher developed the theory of junctions for full sibmating, and the theory of junctions was quickly extended towards self-fertilization (Bennett 1953), alternate parent-offspring mating (Fisher 1959; Gale 1964), random mating (Stam 1980) and recombinant inbred lines (Martin and Hospital 2011). In order to derive expected numbers of junctions and variation in the number of junctions, Fisher had to assume that the size of the genetic blocks delineated by these junctions was exponentially distributed. Using simulations, Chapman and Thompson (2003) showed that this assumption was inaccurate, and that large blocks tended to be overrepresented compared to an exponential block size distribution. Furthermore, Chapman and Thompson extended the theory of junctions towards populations growing in size at a constant rate, and towards subdividing populations (Chapman and Thompson 2002, 2003).

Within the theory of junctions, the chromosome is assumed to be continuous, and to be infinitely divisible. Given that a chromosome consists of an array of base pairs, such an assumption provides an accurate approximation only if the number of base pairs is extremely large. However, ancestry data is usually not acquired on the level of base pairs, but rather using a limited number of markers (microsatellites, single nucleotide polymorphisms (SNPs), single feature polymorphisms (SFPs)) per chromosome. Unfortunately, the number of markers required to detect all haplotype blocks needs to be high; in order to detect 90% of all apparent haplotype blocks at least 10 times more markers than blocks are required (MacLeod *et al.* 2005). One way to circumvent such high marker numbers is by comparing simulated haplotype block densities with observed haplotype block densities and apply a post-hoc correction to the molecular data (Buerkle and Rieseberg 2008). Both using extremely high marker densities and performing post-hoc corrections are ad hoc solutions, and both these approaches lack a solid theoretical underpinning connecting current standing theory for the number of blocks in continuous chromosomes, with theory describing the number of blocks in chromosomes described by discrete numbers of markers.

Furthermore, the theory of junctions has focused severely on the idealized situation in which the genome of all F1 hybrid individuals contains equal proportions of the genetic material of the parental species. In nature, the overall ancestry contribution of parental species to the founding hybrid swarm may differ (Edmands *et al.* 2005; Nolte and Tautz 2010; Stemshorn *et al.* 2011). Deviations from an even ratio can cause the genetic material in the F1 hybrids to sway in favor of one of the ancestral species. Strong deviations from equality can increase the impact of drift, as fixation of genetic material in a population becomes more likely, especially for small population sizes. The impact of deviations from an even ancestry contribution of both ancestral species and the interaction with population size remains understudied so far.

Here we present a universal haplotype block theory describing the mean number of haplotype blocks in the population. Our universal haplotype block theory includes both continuous and discrete chromosomes, it takes into account the ancestry distribution of the parental species in the founding hybrid swarm, and it includes drift due to finite population size. We compare our theory with previously obtained results by MacLeod *et al.* (2005) and Buerkle and Rieseberg (2008), and confirm validity of our theory using individual based models.

Our paper is structured as follows: first we derive the mean number of haplotype blocks in a continuous chromosome, for infinite and finite population size. Then, we proceed to derive dynamics of the mean number of haplotype blocks for discrete chromosomes consisting of a finite number of recombination sites. We then infer universal haplotype block dynamics, by combining properties of our previous derivations for the mean number of blocks in discrete and continuous chromosomes. Using individual based simulations, we then demonstrate the validity of our derivations, and extend our derivations towards discrete chromosomes in finite populations, and towards multiple recombination events per meiosis.

## 1. Analytical model

### *A*. *The expected number of haplotype blocks in a continuous chromosome*

First, we derive the expected number of blocks depending on the time since the onset of hybridization. We assume infinite population size, random mating, and an continuous chromosome, e.g. there are an infinite number of recombination sites along the chromosome. We assume that only a single crossover event occurs per chromosome per meiosis, which corresponds to the assumption that chromosomes are 1 Morgan long. The recombination rate is assumed to be uniformly distributed across the chromosome. Both chromosomes are interchangeable, and we do not keep track of the identity of chromosomes. Each individual is diploid and chromosomes are inherited independently, which allows us to track haplotype blocks within only one chromosome pair, rather than all pairs simultaneously.

We start by formulating a recurrence equation based on the expected change, or the change in mean number of blocks per generation. Given a recombination site picked randomly across the length of the chromosome, the genomic material on either chromosome can either be identical or different. If the genomic material is identical, no change in the number of blocks occurs. If the genomic material is different, a new block is formed (see Figure 1). The probability of observing the same type of genomic material on both chromosomes is proportional to the frequency of genomic material of that type in the population, which in turn is dependent on the frequency of the corresponding ancestral species in the ancestral population. We denote the frequency of type *P* genomic material from ancestral species 1 as *p*, and the frequency of genomic material of the other type *Q* (from the other ancestral species) as *q*, where *p* = 1 − *q*. The probability of having the same type of genomic material on both chromosomes at the recombination site is then *p*^2^ + *q*^2^, in which case no change in the number of blocks is observed. With probability 2*pq* the type of genomic material on both chromosomes differs and an increase in number of blocks is observed. We obtain

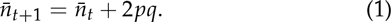

Here *n̅_t_* is the average number of blocks at time *t*. The solution of Equation (1) is given by

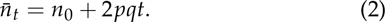

The number of blocks increases linear in time. The probability of having a different type of genomic material, 2*pq*, is the heterozygosity *H*, and we can write Equation (2) in terms of heterozy *n̅_t_* = *n*_0_ + *Ht*. Taking into account that the number of junctions *J* at time *t* is *J_t_ = n_t_* − 1 and assuming a non-constant heterozygosity *H*, we recover the previously obtained result (Chapman and Thompson 2002; MacLeod *et al.* 2005; Buerkle and Rieseberg 2008)

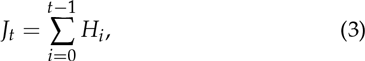

where *J_t_* is the number of junctions *J* at time *t*. Because the population size is infinite, in our case the average heterozygosity does not change from *H*_0_, and *J_t_* = *H*_0_*t*.

If the population size is not infinite, but finite with size *N_j_* at time *j*, and we allow for selfing, the average heterozygosity changes over time (Crow and Kimura 1970) as

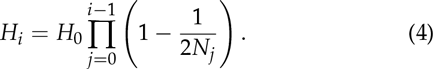

Assuming constant population size, *N_j_* = *N* for all *j*, the expected number of junctions is given by

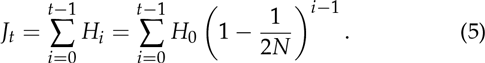

The expected number of blocks, given an initial proportion *p* of species 1 is given by:

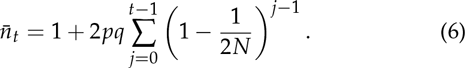

In the limit of *N* → ∞ to we recover Equation (2). For *t* → ∞ to, we have (MacLeod *et al.* 2005)

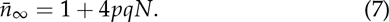

Thus, for finite *N*, the number of blocks converges to a finite number determined only by the population size and the initial frequencies of genomic material, which in turn depends on the frequency of the ancestral species during the first admixture event.

**Figure 1.**
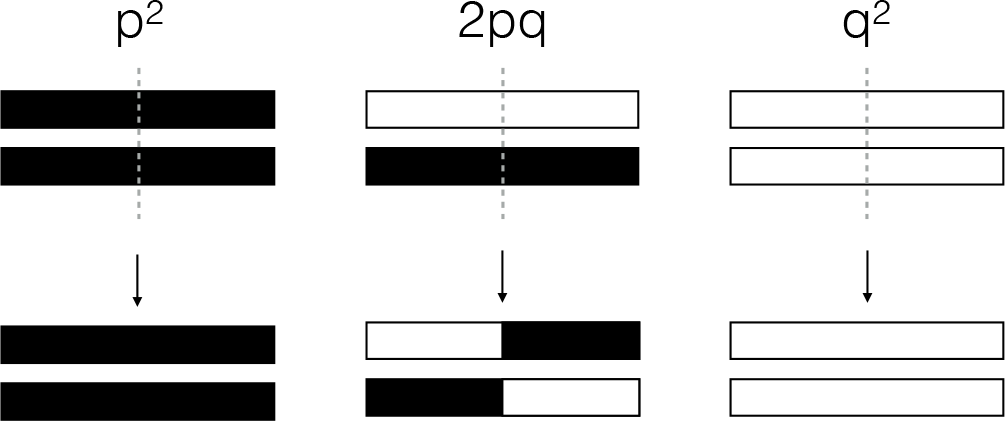
Change in number of haplotype blocks depending on the genomic match between blocks. Top row depicts the two parental chromosomes, bottom row depicts the two chromosomes produced after recombination has taken place at the grey dotted line during meiosis. Genome types are indicated using either black color (type *P*), or white color (type *Q*).With probability *p*^2^ + *q*^2^ no change in the number of blocks is observed. With probability 2*pq* we observe an increase in the number of blocks, where *p* = 1 − *q* is the fraction of genomic material of type 1.

### *B*. *A finite number of recombination spots: a discrete chromosome*

In the previous section we have assumed that recombination never occurs twice at the same spot. In reality, a chromosome can not be indefinitely divided into smaller parts. We therefore proceed to study the change in number of blocks in a chromosome consisting of *L* different chromosomal segments, where each segment represents a minimal genomic element that can not be broken down further, for instance a single nucleotide, a gene, a specific codon, or a genomic area delineated by two genetic markers. Considering a chromosome of *L* genomic segments, there are *L* − 1 possible crossover spots (junctions). Given that there are *n* blocks on the chromosome, there are *n* − 1 points on the chromosome where one block ends, and a new block begins (Fisher called these external junctions), excluding the tips of the chromosome. The probability that a recombination event takes place at exactly such a point, given that there are *n_t_* blocks at time *t* is:

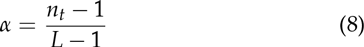

Conditioning on the type of the first chromosome, and only looking at the first of the two produced chromosomes (all other produced chromosomes are identical or the exact mirror image of this chromosome) we can distinguish four possible events, taking into account the location of the recombination spot on both chromosomes (Figure 2):

A) recombination takes place on an existing junction on both chromosomes (probability *α*^2^)
B) recombination takes place on an existing junction on one chromosome, and within a block on the other chromosome (probability *α*(1 − *α*)
C) recombination takes place on within a block on one chromosome, and on an existing junction on the other chromosome (probability (1 − *α*)*α*)
D) recombination takes place within a block on both chromosomes (probability (1 − *α*)^2^)

**Figure 2.**
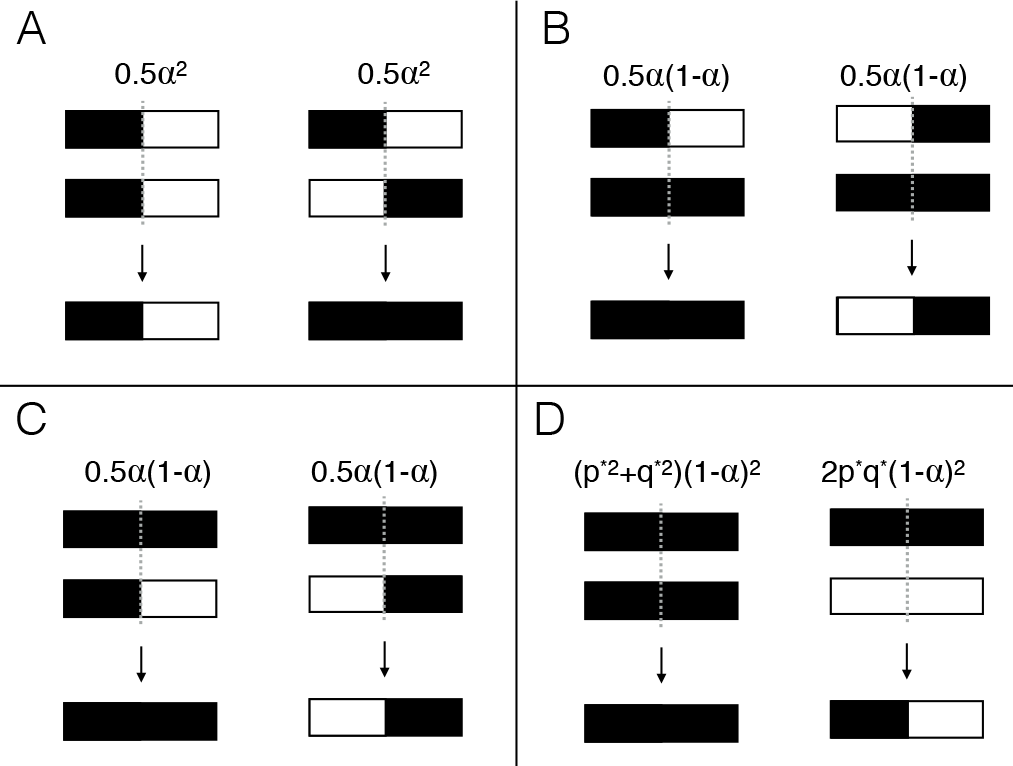
Change in number of haplotype blocks depending on the genomic match between blocks. Full chromosomes are shown here, but the same rationale applies to subsets of a chromosome. Top rows within each panel indicate the two parental chromosomes, bottom row indicates one of two possible resulting chromosomes after meiosis, where recombination takes place at the dotted grey line. Genomic material of type 1 is indicated in black, genomic material of type 2 is indicated in white. With probability *α*^2^ recombination takes place on an existing junction on both chromosomes **A**, with probability *α*(1 − *α*) recombination takes place on an existing junction on one chromosome, and within a block on the other chromosome **B**, with probability (1 − *α*)*α* recombination takes place on within a block on one chromosome, and on an existing junction on the other chromosome **C**, with probability (1 − *α*)^2^ recombination takes place within a block on both chromosomes **D**.

A)When a crossover event takes place on an existing junction on both chromosomes, there is either no change in the numberof blocks (when the two junctions are identical), or a decrease in the number of blocks (when the two junctions are of opposing type). The probability of either event happening is 1/2, yielding an average change in the number of blocks when crossover takes place on an exisiting junction on both chromosomes of −1/2

B) When a crossover event takes place on an existing junction on one chromosome, and within a block on the other chromosome, there are two possibilities: either the block on the other chromosome is of the same type as the genomic material before the existing junction, or it is of the other type. If it is of the same type, the existing junction disappears, and the number of blocks decreases by one. If it is of the other type, the existing junction remains and the number of blocks does not change. The probability of either event happening is 1/2 and hence we expect the total number of blocks on average to change by ‒1/2.

C) When a crossover event takes place within a block on the first chromosome, and on an existing junction on the other chromosome (the inverse of the previous situation), the outcome is exactly the opposite. If the genetic material after the junction on the second chromosome is of the same type as the block on the first chromosome, no new junction is formed and the number of blocks stays the same. If the genetic material after the junction on the second chromosome is of a different type than that of the block on the first chromosome, a new junction is formed and the number of blocks increases by one. The probability of either event happening is 1/2, and hence we expect the total number of blocks on average to change by 1/2.

D) When recombination takes place within a block on both chromosomes, matters proceed as described for the continuous chromosome: with probability *p*^2^ + *q*^2^ we observe no change in the number of blocks, and with probability 2*pq* we observe an increase. But, since we are dealing with a finite number of junction positions along the chromosome, the frequency of junction spots of a genomic type is no longer directly related to *p*. If there would be no blocks, i.e. if the genomic material would be distributed in an uncorrelated way, we know that *p*(*L* − 1) junction spots are of type *P*, that is, they are within a block of type *P*. Similarly, *q*(*L* − 1) junction spots are within a block of type *Q*. As new blocks are formed, the number of junction spots that are still within a block decreases. With the formation of a new block, on average both a junction within a block of type *P* and a junction within a block of type *Q* are lost, such that on average, after the formation of a new junction, the number of junctions of type *P* decreases by 1/2 (*n* − 1). Thus the number of junctions within a block of type *P* is *p*(*L* − 1) − 1/2 (*n* − 1). Similarly, the number of junctions within a block of type *Q* is *q*(*L* ‒ 1) ‒ 1/2 (*n* ‒ 1). The probability then of selecting an internal junction of type *P* is the number of internal junctions of type *P* divided by the total number of junctions. Let us denote the probability of selecting an internal junction of type *P* by *p*\*, which is then given by

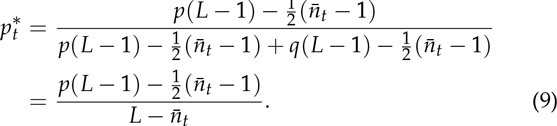

And the probability 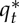 is

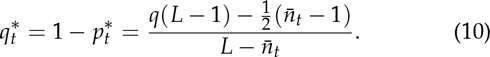

With probability 2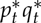 we observe an increase in number of blocks. Combining the scenarios (A)-(D) we can formulate the total expected change in number of blocks

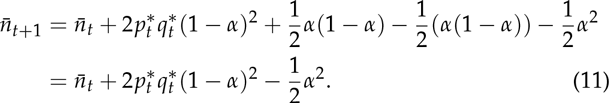

In terms of *p, q*, and *L*, Equation (11) can be written as

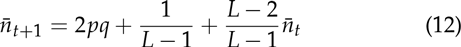

The solution of the recursion Equation (12) is given by

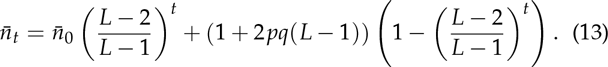

The exponential decay terms ensures that we have convergence at *t* → ∞, where we obtain

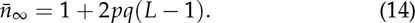

A Taylor expansion at *t* = 0 shows that initially, the number of blocks increases linearly

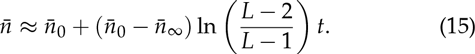

### *C*. *Multiple recombination events*

So far, we have assumed that during meiosis only a single crossover event occurs. Although this might often apply, multiple crossover events occur frequently. First, we consider the case of two crossover events. Assuming that the position of the two crossovers is independent, that there is no interference between the two crossovers, and that the two crossovers do not take place at the same position, we can extend our recurrence equations as follows.

For an infinite population, with a discrete chromosome, the first position is still chosen as in Equation (11). To obtain the probability for the second position of selecting a junction that lies within two dissimilar blocks, we have to correct 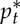 and 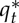 (whereas previously there were *L* − 1 spots, there are now *L* − 2), and we obtain

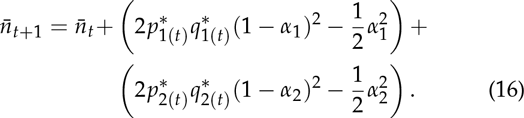

Where:

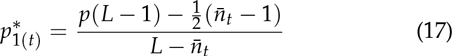

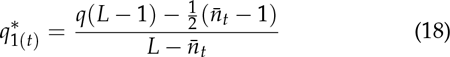

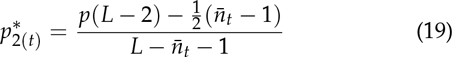

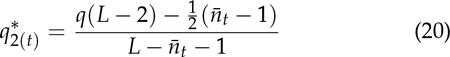

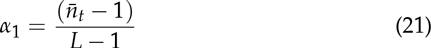

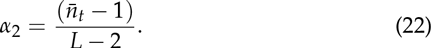

Equation (16) has the solution:

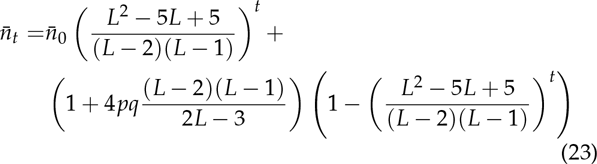

Again we have convergence at *t* → ∞ where we obtain:

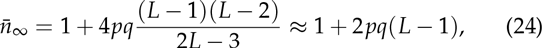

where the approximation holds for large *L*. The difference in the maximum number of blocks between one and two recombination events is

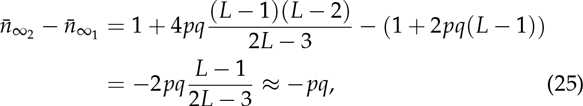

where the approximation again holds for large *L*. An increase in the number of recombination events thus decreases the maximum number of blocks.

Extending Equation (16) towards *M* recombination events is similar,

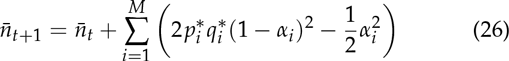

with

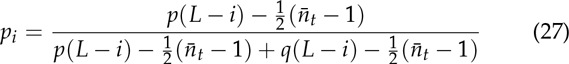

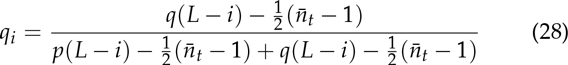

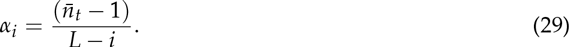

For not too small *L*, and not too large *M*, taking into account Equation (25), we can approximate *n̅*_∞_ by:

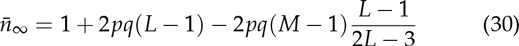

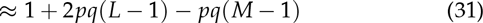

Using numerical iteration of Equation (26) for *M* = [2, 3, 4, 5] and *L* = [2*M* + 1, 2*M* + 2, … , 200], and comparing the maximum number of blocks with the approximation of Equation (31) shows that Equation (31) is a good approximation (error < 0.1%) for *L* ≥ 10*M* − 3.

## 2. Universal haplotype dynamics

The general pattern of the accumulation of blocks over time is highly similar across different scenarios: after an initial period of a strong increase in the number of blocks, the number of blocks starts to increase more slowly, and approaches a maximum. Generally, we can infer that the maximum number of blocks is dependent on *p*, *N* and *L*, as do the time dynamics required to reach this maximum. Hence, we can describe the number of blocks relative to the number of blocks at *t* = ∞, which we define here as *K*. During meiosis, the average number of blocks in the most ideal case can increase with *γ* blocks. There might be limitations, dependent on *p, N, L* or the number of already existing blocks. As such we make the ansatz:

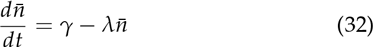

where *γ* is the maximum growth rate and *λ* encompasses all factors limiting the formation of new blocks. This includes, but is not limited to, factors induced by a finite population size *N* by a finite chromosome size *L*. Defining *τ* = *λt*, we can rescale time in Equation (32) such that

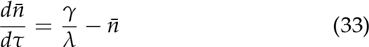

Measuring the number of blocks in terms of their equilibruim values, 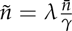, we obtain

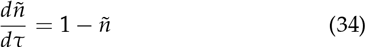

The solution of Equation (34) is of the form:

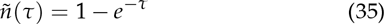

Given that at *τ* = 0, a chromosome, by definition, consists of one block, the number of blocks at *τ* = 0 has to be equal to 1/*K*: *ñ*(0) = 1/*K*. Thus, in order to accurately describe haplotype block dynamics, we expect the dynamics to be of the form

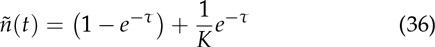

We can find the scalaing of time by solving

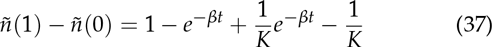

By definition *ñ*(0) is always 1/K, which leads to

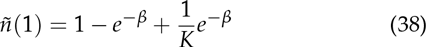

This allows us to calculate *β* from *K* and *ñ*(1),

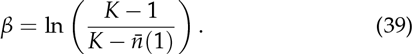

Returning to our original notation, the haplotype block dynamics is geven by:

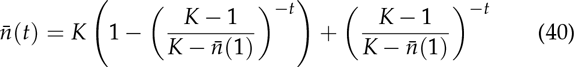

For *t* → ∞, this converges to *K*, and for *t* = 0 is equal to 1. Equation (40) provides us with a general scalable equation where all haplotype block dynamics are described in terms of *K* and *n̅*(1).

### *A*. *Implementing universal dynamics*

To implement the universal equation (40), we only require to know *K* and *n̅*(1). For an infinite population, with a continuous chromosome, we have previously derived that the average number of blocks at time *t* is given by: *n̅*_t_ = 1 + 2*pqt* (equation 2), and thus *n̅*_t=1_ = 1 + 2*pq*. The maximum number of blocks *K* is equal to ∞.

For a finite population with a continuous chromosome, we have shown that *K* = 1 + 4*pqN* and that the number of blocks at time *t* is given by 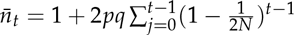 (Equation (6)), and hence *n̅*_*t*=1_ = 1 + 2*pq*.

**Figure 3.**
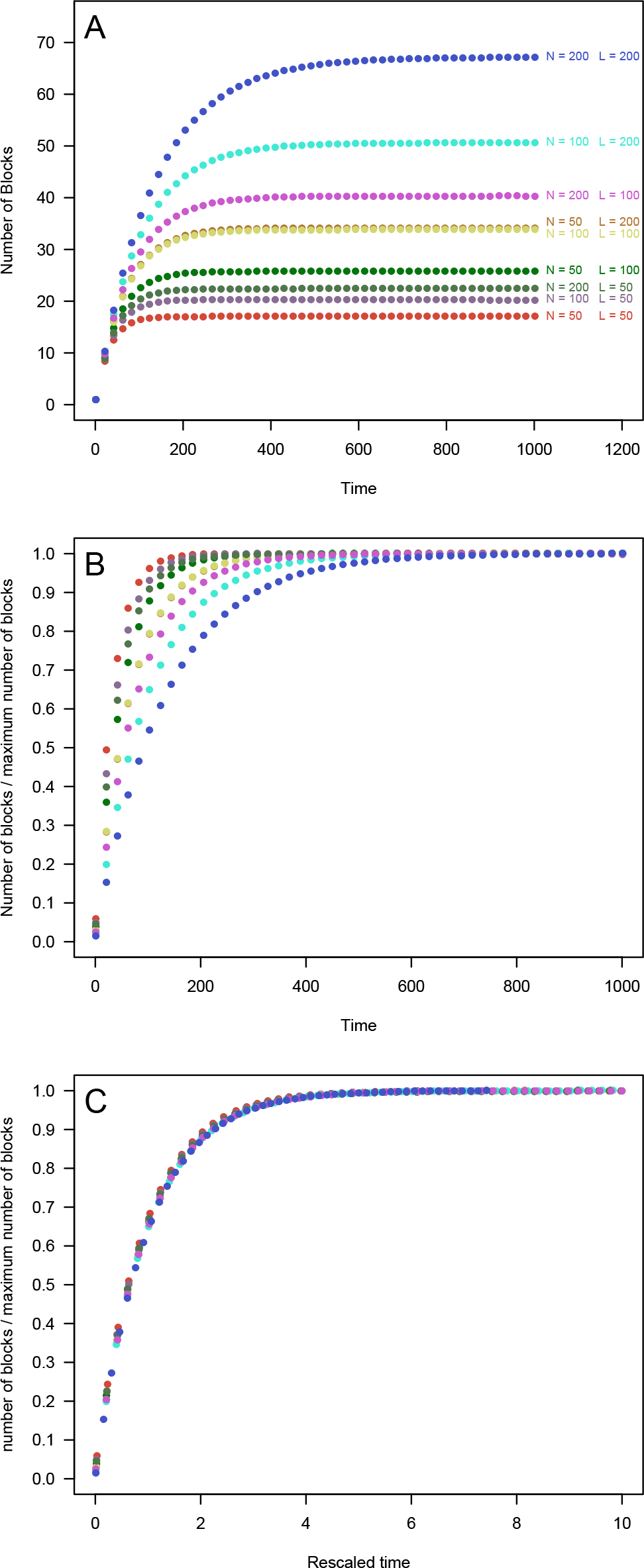
Graphical example of the construction of universal haplotype block dynamics using results from individual based simulations. **A**: mean number of haplotype blocks for *p* = 0.5, *N* = [50,100,200] and *L* = [50,100,200], number of replicates = 10,000. **B**: The mean number of blocks for the same parameter combinations, after rescaling the number of blocks relative to the maximum number of blocks *K*. **C**: The rescaled number of blocks vs rescaled time, by rescaling time according to *β* in Equation (39). After rescaling both the number of blocks according to *K*, and time according to *β*, all curves for different values of *N* and *L* reduce to a single, universal, curve, which follows Equation (40).

For an infinite population with a discrete chromosome, we have shown that *K* = 2*pq*(*L* − 1) + 1 and that the number of blocks at time *t* is given by 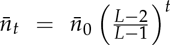 + 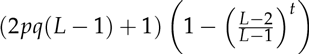 (Equation (13)), and hence:

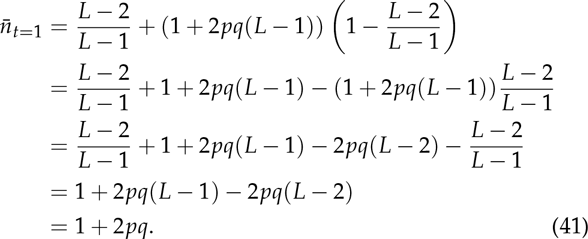

We find that regardless whether the chromosome is continuous or discrete, and regardless of whether the population is finite on infinite, *n̅*(1) = 1 + 2 *pq,* which makes intuitive sense: in the first generation, none of the factors that limit recombination as a result of finite population size, or finite chromosome size come into play. When the population is finite, the formation of new blocks is limited by recombination taking place at a recombination spot where in a previous generation recombination has already taken place. In the first generation, all chromosomes are non-recombined, and finite population effects have no effect yet. When the chromosome is discrete, the formation of new blocks is limited by recombination taking place on a site that has previously recombined. In the first generation, no previous recombination events have happened yet, and recombination is thus not limited (yet).

## 3. Individual based simulations

To verify our analytical framework, and extend the framework towards discrete chromosomes in finite populations, we test our findings using an individual based model. We model the hybrid population as a Wright-Fisher process, extended with recombination:

- Non-overlapping generations
- Constant population size *N*
- Random mating
- Diploid
- Uniform recombination rate across the genome
- *M* recombination events per meiosis

Each individual has 2 chromosomes of length *L*, which are a sequence of 0 and 1’s, where 1 represents an allele from an ancestral parent of type *P* and a 0 represents and allele from an ancestral parent of type *Q*. The model works as follows. In the first time step, *N* individuals are generated, where each individual can have either two parents of type *P* (with probability *p*^2^), two parents of type *Q* (with probability (1 − *p*)^2^) or one parent of type *P* and one parent of type *Q* (with probability 2*p*(1 − *p*)).

In every consecutive time step, *N* new individuals are produced, where each individual is the product of a reproduction event between two individuals (including selfing) from the previous generation. Parental individuals are drawn with replacement, such that one individual could reproduce multiple times, but will on average reproduce one time. We assume that in a mating event both parents produce a large number of haploid gametes from which two gametes (one from each parent) are chosen to form the new offspring. During production of the gametes, *M* recombination sites are chosen. The location of the recombination sites follows a uniform distribution between 0 and *L*.

### *A*. *Continuous Chromosome*

We model each chromosome as a continuous line, and only keep track of junctions delineating the end of a block. For each junction, we record the position along the chromosome (a number between 0 and 1) and whether the transition is 0 → 1 or 1 → 0. Over time, the number of blocks reaches a maximum value, but only if the population is finite (Figure 4). In the first few generations the accumulation of blocks follows that of an infinite population (dotted line in Figure 4), but rapidly simulation results start deviating from the infinite population dynamics. The maximum number of blocks is roughly obtained within 10*N* generations. Furthermore, when the amount of genetic material from either of the ancestral species is strongly skewed (e.g. *p* = 0.9), the maximum number of blocks is lower, and is reached within a shorter timespan.

### *B*. *Discrete Chromosome*

To mimic a chromosome consisting of a discrete number of genomic elements, we model the chromosome as a bitstring, where a 0 indicates a chromosomal segment of type 0, and a 1 indicates a chromosomal segment of type 1. To approximate an infinite population we use a population of size 100,000. Mean number of blocks in the stochastic simulations closely follow our analytical estimates (Figure 5), for all chromosome lengths considered here. The maximum number of blocks is reached in roughly 10*L* generations. Again we observe that for strongly skewed ancestral proportions (*p* = 0.9), the maximum number of blocks is lower, and the maximum is reached within a shorter timeframe.

### *C*. *Discrete Chromosome in a Finite Population*

We have shown in section 2 that we can describe haplotype dynamics for a population of any *N* and a chromosome of any *L*, as long as either *N* or *L*, or both are infinite. To describe haplotype dynamics for any given *N* and *L*, we only need to know *K* and *n̅*(1). Since *n̅*(l) = 1 + 2*pq* for any *N* and *L*, this leaves us with disentangling the relationship between *N, L* and *K*. We know that for a given *L*, with varying *N, K* will approach 2*pq*(*L* − 1) + 1 when *N* → ∞ (Figure 6A). The exact functional relationship however, remains unknown. Similarly for given *N*, with *L* → ∞, *K* will approach 4*pqN* + 1 (Figure 6B). But again, the exact functional relationship remains unknown. We therefore turn to individual based simulations to generate values of *K* for different combinations of *L* and *N*. Then, using non-linear least-squares fitting we formulate an approximate functional relationship between *N, L*, and *K*. Nevertheless, such a formulation will remain a crude approximation only and should be interpreted as such. In order to do so, we simulate 10,000 populations for all combinations of *N* and *L* for values [10, 20, 30,… ,200, 250,300,…, 500] and *p* = [0.5,0.6,0.7,0.8,0.9]. We then use the mean of the maximum number of haplotype blocks across these 10,000 replicates to reconstruct our functional relationship. Given a value of *L*, we guess the relationship between *N* and *K* to be approximately of the form: *K_N_* = (2*pq*(*L* − 1) + 1)*N*/(*aN* + *b*)), where *a* and *b* are to be determined and are possibly dependent on *L* and *p*. Note that we could have also chosen another form, such as (2*pq*(*L* − 1) + 1)(1 − 1/(*aN*)^*b*^) or (2*pq*(*L* − 1) + 1)(1 − exp(− *aN* + *b*)), however, we found *K_N_* = (2*pq*(*L* − 1) + 1)*N*/(*aN* + *b*)), to be easiest to fit to the data. Using non-linear least squares estimation we find that for *L* = 100 and *p* = 0.5, *a* = 1.014 and *b* = 47.83 (Figure 6 A). We can repeat this process for different values of *L* (but keeping *p* = 0.5) and find that *a* is always close to 1 and that *b* is always close to 2*pq*(*L* − 1) + 1. Hence, it makes more sense to find our approximation in the form: *K_N_* = *c*(*2pq*(*L* − 1) + 1 )*N*/(*aN* + (2*pq*(*L* − 1) + 1))) (fitting more than 2 parameters at the same time tends to lead to inaccurate results). We find, by fitting to varying values of *N* and *p*, that *a* and *c* both tend to 4*pq*. With that, we obtain the approximation for *K*:

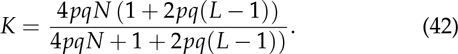

**Figure 4.**
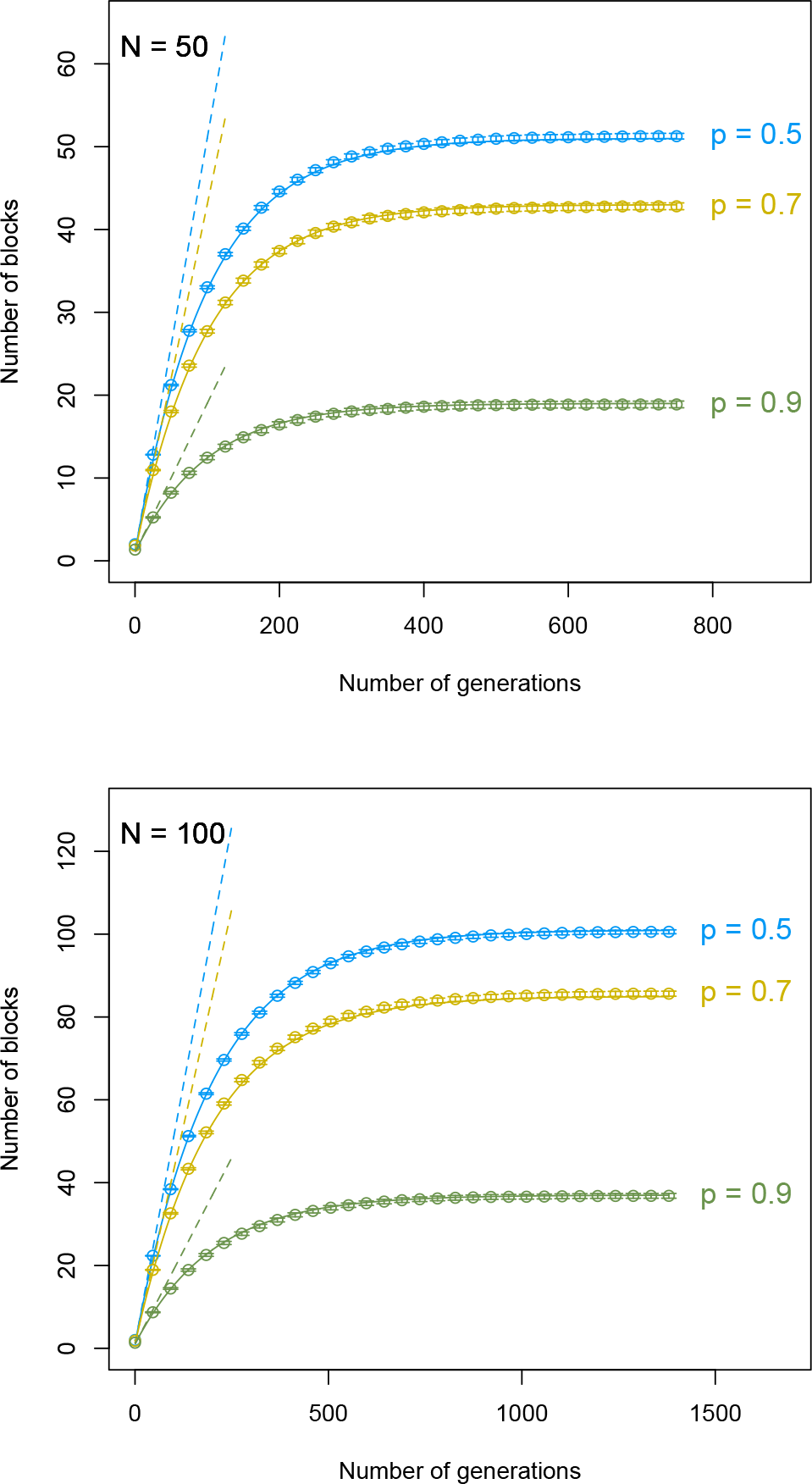
Number of haplotype blocks over time for stochastic simulations assuming a continuous chromosome and a population size of 50,100 individuals (circles), the analytical prediction for an infinite population size (dotted line), or the analytical prediction for a finite population size (Equation (40), *K* = 4*pqN* + 1), solid line. Error bars indicate the standard error of the mean across 1,000 replicates. Shown are results for different initial frequencies of the two parental species *p,* such that the initial heterozygosity *H*_0_ = 2*p*(l − *p*).

**Figure 5.**
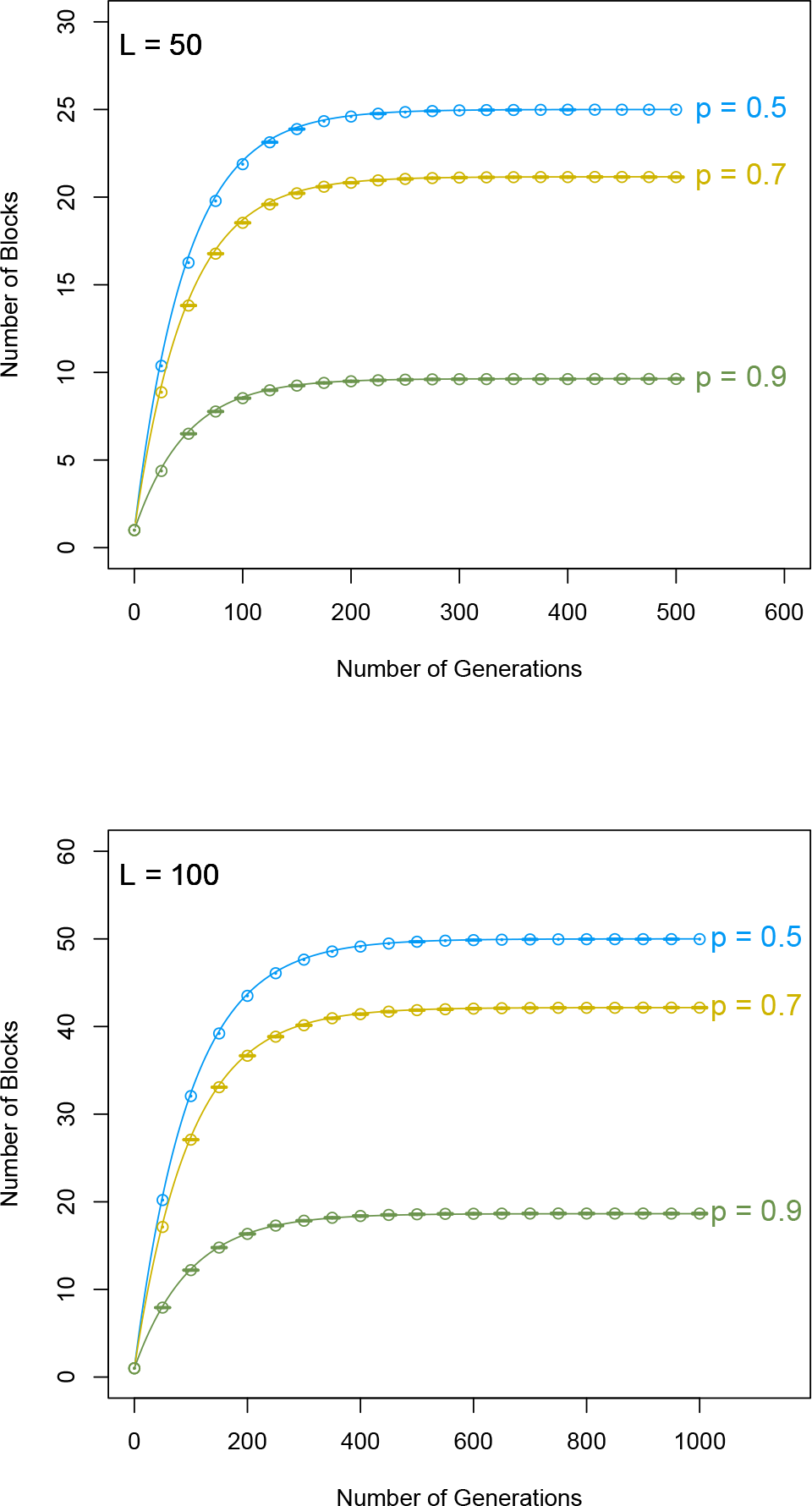
Number of haplotype blocks over time for either stochastic simulations assuming a continuous chromosome and a population size of 100,000 (circles), or the analytical prediction according to Equation (40), *K* = 2*pq*(*L* − 1) + 1 (solid line). Error bars indicate the standard error of the mean across 100 replicates. Shown are results for different initial frequencies of the two parental species *p*, such that the initial heterozygosity *H*_0_ = 2*p*(l − *p*).

We recover both limits of *K* for *N* → ∞ and *L* → ∞, and formulate our general approximation as:

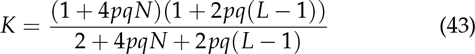

Comparing values of *K* expected following Equation (43) with the observed mean estimates from the simulations confirms that our approximation provides estimates that are close to the mean estimates (Figure 6C, Observed vs Expected, intercept = −0.0727, slope = 0.9987, *R*^2^ = 0.9999). Furthermore, equation (43) reduces to Equation (7) when *L* ≫ *N*, and reduces to Equation (14) when *N* ≫ *L*. As such, Equation (43), albeit an approximation for *K*, encompasses all combinations of *N* and *L*. To extend Equation (43) towards *M* recombinations per meiosis, e.g. to chromosomes of any maplength in Morgan, we can suffice with a more sparse grid. Here we only need to show (numerically) that we can correct *K* for *M* recombinations, and substitute the corresponding equations into Equation (43). It has been shown previously that *K*, for *L* → ∞ is given by: *K* = 4*pqMN* + 1 (Buerkle and Rieseberg 2008) and *K* for *N* → ∞ is given by: *K* = 2*pq(L* − 1) + 1 − *pq*(*M* − 1) (Equation (31)). Substituting these expressions for *K* into Equation (43) we obtain

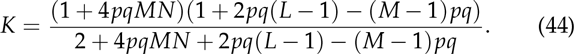

We simulate values of *K*for combinations of *N* and *L* in [50, 100, 150, 200], *p* in [0.5, 0.9] and *M* in [2,3,4]. We find that results from individual based simulations are very close to our predicted equation (Figure 6, intercept = −0.1352, slope = 1.0010, *R*^2^ = 0.9999). Equation (44), just like Equation (43), reduces to *K* = 4*pqMN* + 1 for *L* ≫ *N*, and reduces to *K* = 2*pqM*(*L* − 1) − 1 − *pq*(*M* − 1) for *N* ≫ *L*.

As a further test of the accuracy of Equation (44), we repeat simulating values of *K*, but for more typical maplengths found in molecular data *M* = [0.5, 0.75, 1, 1.25, 1.5, 1.75, 2] (using the same sparse grid for *N, L* and *p*). We interpret fractional numbers of recombinations as a mean rate, such that an average of 1.25 recombinations implies that in 25% of all meiosis events there are 2 recombinations, and in 75% of all meiosis events there is 1 recombination. Again we find that obtained mean estimates for *K* are close to those predicted using Equation (44) (intercept = −0.2618, slope = 1.0027, *R*^2^ = 0.9999, not shown in Figure 6).

**Figure 6.**
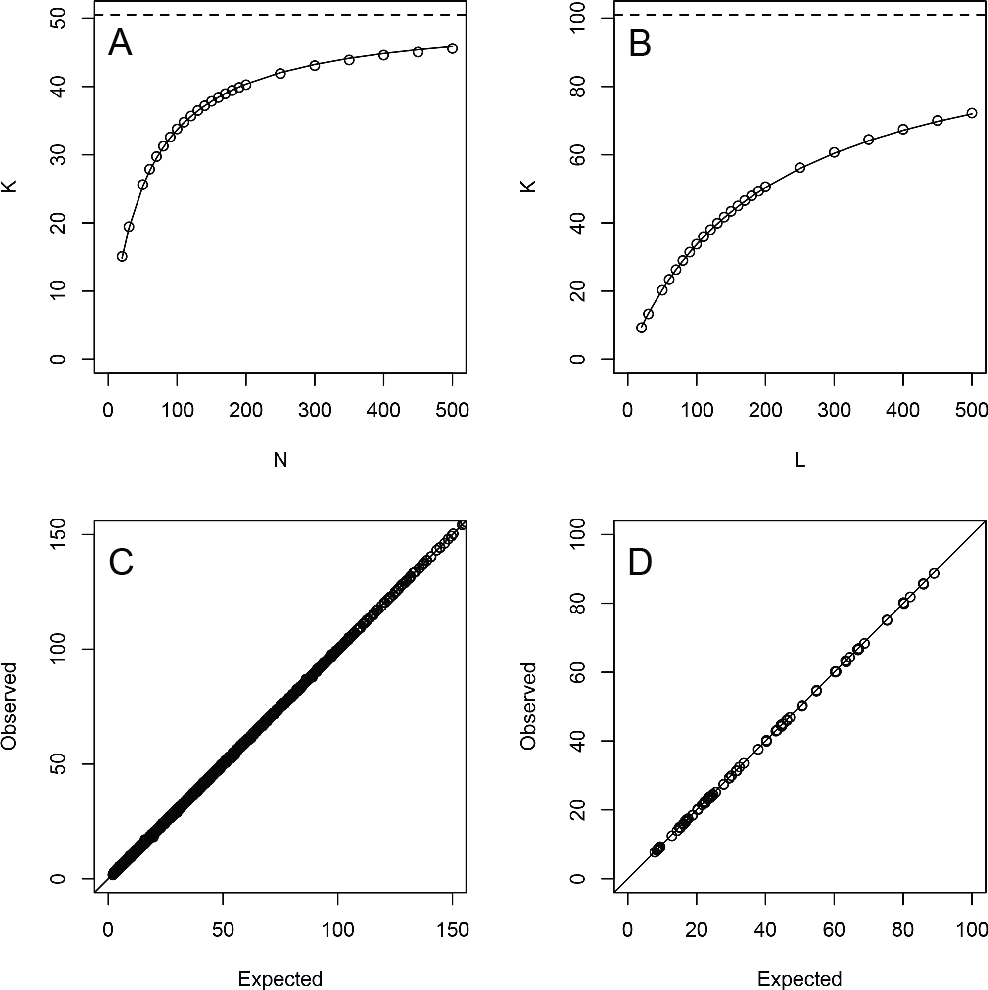
Numerical derivation of *K* for a finite population and discrete chromosome. **A**: Example relationship between *K* and *N*, for *L* = 100 and *p* = 0.5. Points are the mean of 10,000 individual based simulations. The solid line is the expected value of *K* following Equation (43). The dotted line represents the upper limit of *K* for *N* = to. **B**: Example relationship between *K* and *L*, for *N* = 100 and *p* = 0.5. Points are the mean of 10,000 individual based simulations. The solid line is the expected value of *K* following Equation (43). The dotted line represents the upper limit of *K* for *L* = ∞. **C**: Observed vs expected values for *K* for one recombination per meiosis for all combinations of *N* and *L* in [10, 20, 30 … 200, 250, 300 … 500], and *p* in [0.5, 0.6, 0.7, 0.8, 0.9] (3380 combinations, 10,000 replicates per combination). Expected values were calculated using Equation (43). *R*^2^ = 0.9999, slope = 0.999. **D**: Observed vs expected values for *K* for *M* recombinations per meiosis, for all combinations of *N* and *L* in [50,100,150, 200], *p* in [0.5 ,0.9] and *M* in [2,3,4] (96 combinations, 10,000 replicates per combination). Expected values for *K* were calculated using Equation (44). *R*^2^ = 0.9999, slope = 1.0010.

## 4. Discussion

We have shown here how the number of haplotype blocks changes over time after a hybridization event. We have obtained analytical expressions for the change over time in haplotype blocks for discrete chromosomes, in infinite populations or in finite populations (including random drift) and for different frequencies of both ancestral species. Furthermore, we have developed a unifying framework that describes haplotype block dynamics for both discrete and continuous chromosomes of arbitrary map length, for any population size and for any initial frequency of both ancestral species.

We have extended standing theory for haplotype block dynamics for chromosomes with an infinite number of recombination spots, towards chromosomes that consist of discrete genetic elements with a finite number of recombination spots. Molecular data describing these elements typically presents itself in the form of genetic markers (microsatellites, SNPs, (SFP)) that can be traced back to either of the two parental species. The genomic stretches delineated by these markers then form the genetic elements we have considered here, although in principle our framework also applies on the nucleotide level. We have assumed that the genetic elements are of equal size, and are uniformly distributed. Data based on molecular markers often deviates from this assumption, with markers non-uniformly distributed across the genome and the genomic stretches delineated by these markers differing in size. Deviations from a uniform distribution of markers across the chromosome could potentially lead to an underestimation of the number of blocks, as small blocks are more likely to remain undetected.

The universal haplotype block dynamics framework we have presented here provides a neutral expectation for the number of blocks over time. There is a wide range of processes that can cause deviations from our framework, which fall into two distinct categories: either processes acting upon the underlying genomic content, or processes that affect population dynamics. An important process acting upon the underlying genomic content is selection. If a genomic region is under selection, we expect to find a high frequency of genomic material from the beneficial parent. This reduces recombination events in this region, as recombined individuals are selected against (Kimura 1956; Lewontin and Hull 1967).As a result, we expect this region to be homozygous more often, and to have a lower potential for the formation of new blocks. We thus expect selection for either of the parental types to slow down the formation of haplotype blocks. Future work could focus on the minimal level of selection to offset neutral haplotype block dynamics or whether deviations from neutral haplotype block dynamics can be used to identify genomic areas that are under selection.

Alternatively, the formation of new blocks could speed up when the combination of alleles from both parents provides a selective advantage. The resulting overdominance favours a heterozygous genotype (Maruyama and Nei 1981)), in turn favouring the formation of new blocks (equations (2), (6) and (13)). Likewise, positive epistasis among linked loci could favour a combination of alleles of different parents. As a result, recombination between these loci would be selected for, speeding up block formation. However, such increases in speed would only be seen as short bursts, because after establishment or fixation of favourable alleles, block formation is no longer affected.

A non-uniform recombination rate across the chromosome could cause deviations from neutral haplotype block dynamics. Empirical work has shown that recombination rates are often not equal across the chromosome, but are increased towards the peripheral ends of the chromosome (Lukaszewski and Curtis 1993; Pan *et al.* 2012; Roesti *et al.* 2012). As long as the chromosome is continuous, the exact shape of the recombination landscape has no effect on haplotype block dynamics. For a discrete chromosome however, haplotype block dynamics change (see AppendixA for a demonstration of increased recombination towards the peripheral ends). If some sites have an increased probability of recombination compared to others, recombination is more likely to occur at a site that has already experienced a recombination event before. As a result, the formation of new haplotype blocks is slower than expected under uniform recombination. The maximum number of blocks that can be reached remains unaffected. Similarly, the presence of recombination hotspots, areas in the chromosome with an elevated recombination rate (Gerton *et al.* 2000; Myers *et al.* 2005; Smagulova *et al.* 2011), also slows down the accumulation of blocks (see Appendix B for a demonstration of the effect of hotspots). Similar to increased recombination rates towards the peripheral ends, hotspots skew the recombination rate distribution to such an extend that recombination is much more likely to take place at a site that has been previously recombined, in which case no new block is formed.

Apart from processes that act upon the genomic content, population level processes are also expected to affect haplotype block dynamics. Firstly, deviations from having a constant-population size over time are expected to cause deviations from our haplotype block dynamics framework. A natural extension of our work would for instance be to include either exponentially or logistically growing populations in order to mimic real life dynamics more closely. In exponentially growing populations, the average heterozygosity does not change (Crow and Kimura 1970), which results in dynamics that resemble an infinite population. Similarly, for logistically growing populations, during the initial growth phase, haplotype block dynamics are expected to closely resemble block dynamics in an infinite population. We do have to take into account that even though the population is growing exponentially, drift effects could interfere and cause deviations from infinite population dynamics (Hallatschek *et al.* 2007). How drift and the rate of growth interact and influence haplotype block dynamics remains the subject of future study. Furthermore, in a growing population, the effect of selection is enhanced (Otto and Whitlock 1997), suggesting important interactions between selection, drift and population dynamics.

Secondly, population subdivision, founder effects, a bottleneck or a permanent decrease in population size could speed up fixation of haplotype blocks in the population through drift. Because haplotype blocks become fixed, the average heterozygosity decreases faster than expected, and the accumulation of new blocks is slowed down. Furthermore, the maximum number of blocks decreases as well (following Equation 43). Depending on the speed and timing of the decrease, individuals in the final population potentially display a larger number of blocks than expected from the current population size, retaining blocks fixed in the population before the population decreased in size.

Thirdly, secondary introgression, where admixture with the parental population after founding the hybrid population takes place, will affect haplotype block dynamics. Secondary introgression introduces new parental chromosomes that have not yet recombined and leads to an apparent reduction of the number of blocks (Pool and Nielsen 2009). The apparent reduction of the mean number of blocks effectively ‘turns back time’. A secondary introgression event introduces haplotype blocks that are disproportionally large, compared to the standing haplotype block size distribution. As such, the haplotype block size distribution can potentially complement the mean number of blocks for inferring processes influencing genomic admixture after hybridization (Pool and Nielsen 2009).

Apart from the before mentioned processes, we expect that there are more processes that can affect the accumulation of blocks, including, but not limited to, sib-mating, interference between recombination events, mutation, segregation distortion and heterochiasmy. Except for overdominance or positive epistasis, all processes mentioned above slow down the accumulation of haplotype blocks. This suggests that the universal framework for haplotype block dynamics that we have presented here provides an upper limit to haplotype block dynamics.

Similar to punctuated admixture events that we have considered here, repeated or continuous gene flow between populations can result in haplotype block structures, where the genomic material of the blocks can be traced back to distinct populations (Payseur and Rieseberg 2016), but where secondary migrants introduce disproportionally large haplotype blocks into the population (Harris and Nielsen 2013; Liang and Nielsen 2014). However, in between migration or phases of increased admixture, the introduced genomic material breaks down into blocks, following similar dynamics as in our framework. Gravel has explored the impact of past migration events on the block size distribution, and was able to use the block size distribution to infer past migration events of human populations from genome data (Gravel 2012). Further studies have extended his approach, and increased the accuracy of inference, and extended his approach towards inferring effective population size and population substructuring (Palamara *et al.* 2012; Harris and Nielsen 2013; Hellenthal *et al.* 2014; Sedghifar *et al.* 2016). These studies rely on simulations in combination with likelihood methods to infer migration events from empirical data. Our framework complements that approach, contributes to a more complete understanding of the processes driving haplotype block dynamics.

We have shown here how the genomic material of two parental species mixes over time after a hybridization event. With the current advances in genomic methods, it is now possible, and affordable, to screen species for recurring haplotype blocks of other, closely related, species. Our framework can then be used inversely, by inferring the time of the hybridization event. Given that there are many processes that can potentially slow down the accumulation of haplotype blocks, inferring the time of hybridization using our universal haplotype block dynamics framework provides the lower time limit, e.g. the minimum age of the hybrid. Caution should be taken however, as our work shows that haplotype block dynamics tend to stabilize relatively quickly (on an evolutionary timescale), where typically the number of blocks reaches a maximum limit in the order of 10*N* or 10*L* generations. As a result, haplotype block patterns are especially useful for recent hybridization events.

## 5. Data availability

Computer code used for the individual based simulations has been made available on GitHub and can be found on: https://github.com/thijsjanzen/Haplotype-Block-Dynamics

## A. Appendix

## *A*. *Non-uniform recombination rate*

In the main text, we have assumed the recombination rate to be uniformly distributed along the chromosome. Such an assumption especially applies to chromosomes mapped in Morgan, but does not apply to chromosomes where we track haplotype blocks in physical distance (basepairs). Typically, the recombination rate across the chromosome changes considerably, where it is not uncommon to recover an increase in the recombination rate towards the peripheral ends of the chromosomes (Lukaszewski and Curtis 1993; Roesti *et al.* 2012). To study the effect of such changes in recombination rate, we have simulated haplotype block dynamics assuming a 10 fold increase in the probability of a crossover event close to the peripheral ends compared to close to the center of the chromosome. A 10 fold increase in recombination rate is in line with empirical studies (Lukaszewski and Curtis 1993; Roesti *et al.* 2012), but should mainly be interpreted as an illustration of the effect of elevated recombination rates towards the peripheral ends, rather than an attempt to accurately mimic empirical patterns. Because crossover events are more likely towards the peripheral ends, the probability of a crossover event taking place at a site that has experienced crossover before increases. As a result we notice that the accumulation of blocks is slower than under the uniform recombination rate predictions (Figure 7). The maximum number of blocks remains unaffected, and different initial ratios between the two ancestral species *p* does not change the general pattern of a slowdown of accumulation of haplotype blocks.

## *B*. *Hotspots*

Non-uniform recombination rates might also manifest themselves as a result of recombination hotspots. Recombination hotspots delineate areas in the genome that recombine more often than other areas, and are well documented in many different species (Gerton *et al.* 2000; Myers *et al.* 2005; Mackiewicz *et al.* 2013; Singhal *et al.* 2015; Arbeithuber *et al.* 2015; Smagulova *et al.* 2011). The density of recombination hotspots is generally large with hotspots occuring every 200kb. Depending on the size of the chromosome, this results in a total number of hotspots per chromosome between 300 and 1200. To demonstrate the effect of hotspots on haplotype block dynamics, we model a chromosome consisting of 1000 genomic elements (*L* = 1000). We assume 100 recombination hotspots scattered randomly across the chromosome, where the recombination rate is 9 times the normal recombination rate, such that 50% of all recombination events take place in recombination hotspots (McVean *et al.* 2004). The locations of the recombination hotspots were determined *apriori* and kept constant across replicates and parameter settings. We find that recombination hotspots slow down the formation of haplotype blocks (Figure 8), as recombination is more likely to take place at a site that has previously recombined, compared to a uniform recombination rate. The maximum number of haplotype blocks remains unaffected.

**Figure 7.**
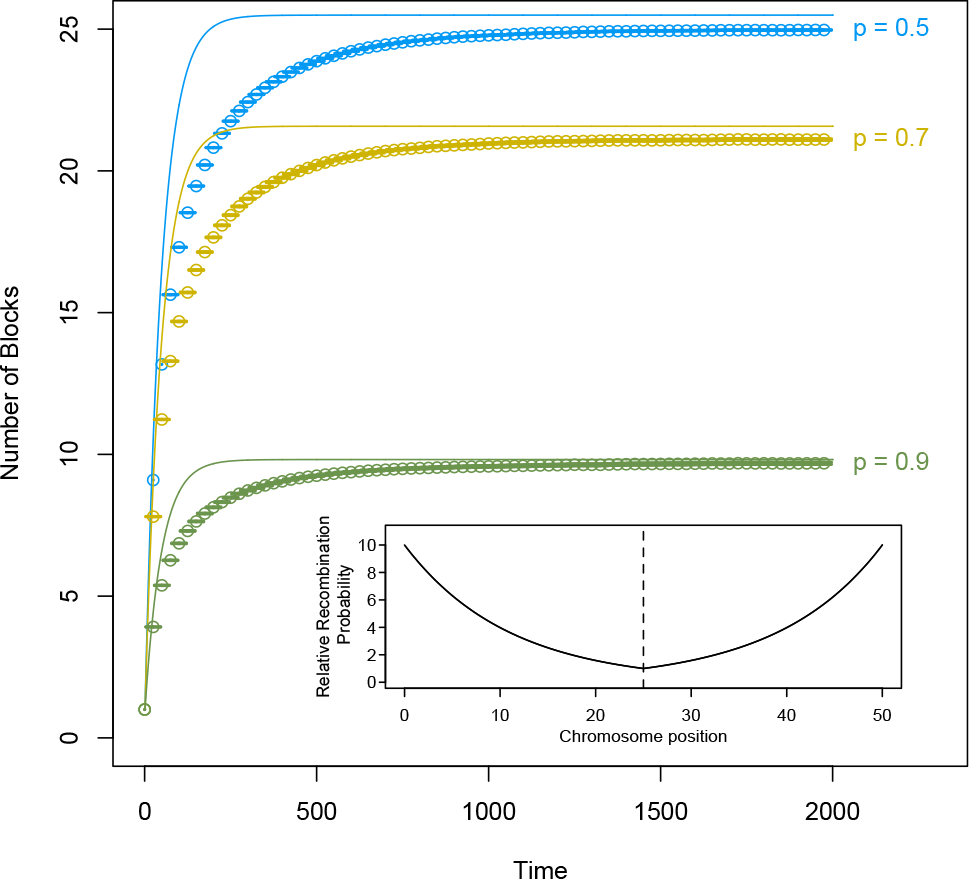
Individual based simulation results for the mean number of blocks over time, assuming an exponential recombination rate distribution with a 10 times higher recombination rate towards the peripheral ends of the chromosome. The recombination rate of site *i* along a chromosome of length *L*, assuming the centromere is located at position *L*/2 is then given by: 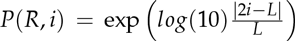. The population size *N* is 100,000 individuals and the number of chromosome elements *L* is 50. The solid line shows the mean number of blocks assuming a uniform recombination rate and infinite population size. Error bars show the standard error of the mean across replicates. The inset graph shows the recombination rate across the chromosome relative to the recombination rate at the centromere, with the dotted line indicating the position of the centromere (at *L*/2).

**Figure 8.**
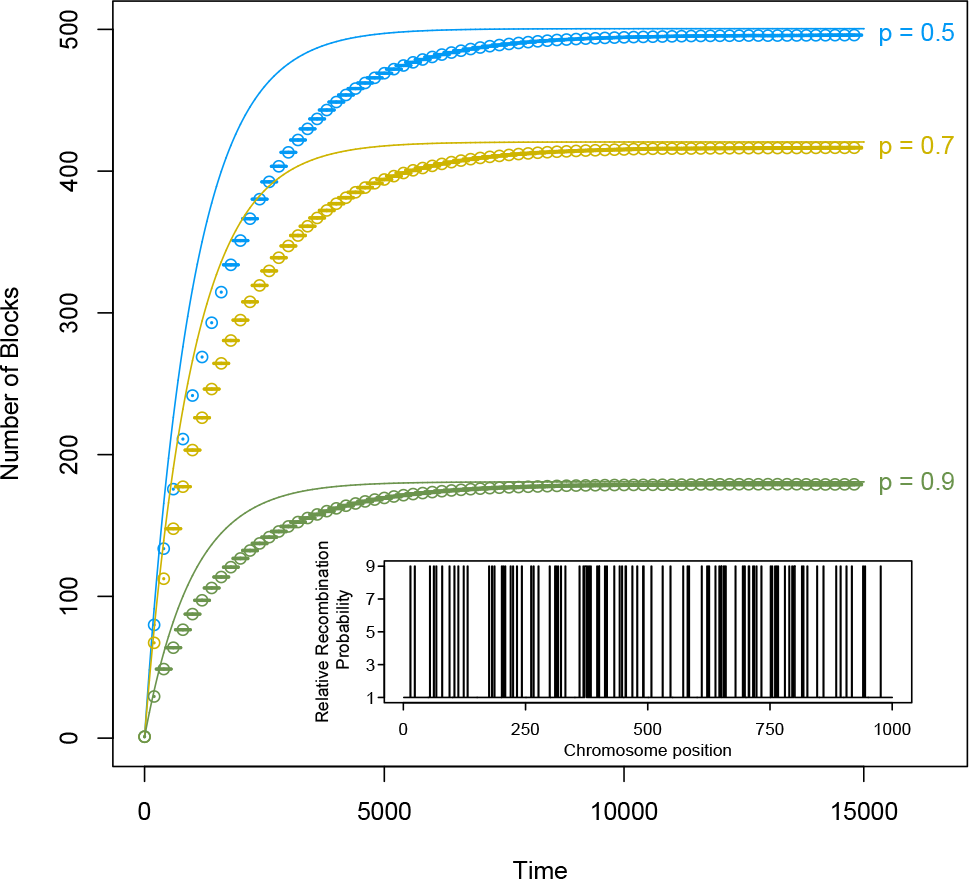
Individual based simulation results for the mean number of blocks over time, assuming 100 randomly placed recombination hotspots on a chromosome consisting of 1000 genomic elements (*L* = 1000), the population size is 100,000. The recombination rate in a hotspot is 9 times the recombination rate outside the hotspots, as a result of which, 50% of all recombination events take place in the recombination hotspots. The dotted line shows the mean number of blocks assuming a uniform recombination rate and infinite population size without hotspots. Error bars show the standard error of the mean across 100 replicates. The inset graph shows the locations of the hotspots across the simulated chromosome.

## Literature Cited

Abbott, R., D. Albach, S. Ansell, J. W. Arntzen, S. J. E. Baird, N. Bierne, J. Boughman, A. Brelsford, C. A. Buerkle, R. Buggs, R. K. Butlin, U. Dieckmann, F. Eroukhmanoff, A. Grill, S. H. Cahan, J. S. Hermansen, G. Hewitt A. G. Hudson, C. Jiggins, J. Jones, B. Keller, T. Marczewski, J. Mallet, P. Martinez-Rodriguez, M. Möst, S. Mullen, R. Nichols, A. W. Nolte, C. Parisod, K. Pfennig, A. M. Rice, M. G. Ritchie, B. Seifert, C. M. Smadja, R. Stelkens, J. M. Szymura, R. Vainola, J. B. W. Wolf, and D. Zinner, 2013 Hybridization and speciation. Journal of Evolutionary Biology 26: 229–246.

Arbeithuber, B., A. J. Betancourt, T. Ebner and I. Tiemann-Boege, 2015 Crossovers are associated with mutation and biased gene conversion at recombination hotspots. Proceedings of the National Academy of Sciences 112: 2109–2114.

Barton, N., 2001 The role of hybridization in evolution. Molecular Ecology 10: 551–568.

Bennett, J., 1953 Junctions in inbreeding. Genetica 26: 392–406.

Buerkle, C. A., R. J. Morris, M. A. Asmussen, and L. H. Rieseberg, 2000 The likelihood of homoploid hybrid speciation. Heredity 84: 441–451.

Buerkle, C. A. and L. H. Rieseberg, 2008 The rate of genome stabilization in homoploid hybrid species. Evolution 62: 266–275.

Chapman, N. and E. Thompson 2003 A model for the length of tracts of identity by descent in finite random mating populations. Theoretical Population Biology 64: 141–150.

Chapman, N. H. and E. A. Thompson, 2002 The effect of population history on the lengths of ancestral chromosome segments. Genetics 162: 449–458.

Crow, J. F. and M. Kimura 1970 An Introduction to Population Genetics Theory. Harper and Row, New York.

Edmands, S., H. Feaman J. Harrison and C. Timmerman 2005 Genetic consequences of many generations of hybridization between divergent copepod populations. Journal of Heredity 96: 114–123.

Fisher, R. A., 1949 The Theory of Inbreeding. Oliver and Boyd.

Fisher, R. A., 1954 A fuller theory of “junctions” in inbreeding. Heredity 8: 187–197.

Fisher, R. A., 1959 An algebraically exact examination of junction formation and transmission in parent-offspring inbreeding. Heredity 13: 179–186.

Gale, J., 1964 Some applications of the theory of junctions. Biometrics pp. 85–117.

Gerton, J. L., J. DeRisi R. Shroff M. Lichten P. O. Brown, and T. D. Petes, 2000 Global mapping of meiotic recombination hotspots and coldspots in the yeast saccharomyces cerevisiae. Proceedings of the National Academy of Sciences 97: 11383–11390.

Grant, V., 1981 Plant speciation. Columbia University Press.

Gravel, S., 2012 Population genetics models of local ancestry. Genetics 191: 607–619.

Hallatschek, O., P. Hersen S. Ramanathan and D. R. Nelson, 2007 Genetic drift at expanding frontiers promotes gene segregation. Proceedings of the National Academy of Sciences 104: 19926–19930.

Harris, K. and R. Nielsen 2013 Inferring demographic history from a spectrum of shared haplotype lengths. PLoS Genetics 9: e1003521.

Hellenthal, G., G. B. Busby, G. Band J. F. Wilson, C. Capelli D. Falush and S. Myers 2014 A genetic atlas of human admixture history. Science 343: 747–751.

Kimura, M., 1956 A model of a genetic system which leads to closer linkage by natural selection. Evolution pp. 278–287.

Lewontin, R. and P. Hull 1967 The interaction of selection and linkage iii synergistic effect of blocks of genes. Der Züchter 37: 93–98.

Liang, M. and R. Nielsen 2014 The lengths of admixture tracts. Genetics 197: 953–967.

Lukaszewski, A. and C. Curtis 1993 Physical distribution of recombination in b-genome chromosomes of tetraploid wheat. Theoretical and Applied Genetics 86: 121–127.

Mackiewicz, D., P. M. C. de Oliveira, S. M. de Oliveira, and S. Cebrat 2013 Distribution of recombination hotspots in the human genome-a comparison of computer simulations with real data. PloS ONE 8: e65272.

MacLeod, A., C. Haley J. Woolliams and P. Stam 2005 Marker densities and the mapping of ancestral junctions. Genetical research 85: 69–79.

Mallet, J., 2007 Hybrid speciation. Nature 446: 279–283.

Martin, O. C. and F. Hospital 2011 Distribution of parental genome blocks in recombinant inbred lines. Genetics 189: 645–654.

Maruyama, T. and M. Nei 1981 Genetic variability maintained by mutation and overdominant selection in finite populations. Genetics 98: 441–459.

McVean, G. A., S. R. Myers, S. Hunt P. Deloukas D. R. Bentley, and P. Donnelly 2004 The fine-scale structure of recombination rate variation in the human genome. Science 304: 581–584.

Myers, S., L. Bottolo C. Freeman G. McVean and P. Donnelly 2005 A fine-scale map of recombination rates and hotspots across the human genome. Science 310: 321–324.

Nolte, A. W., J. Freyhof K. C. Stemshorn, and D. Tautz 2005 An invasive lineage of sculpins, cottus sp. (pisces, teleostei) in the rhine with new habitat adaptations has originated from hybridization between old phylogeographic groups. Proceedings of the Royal Society B 272: 2379–2387.

Nolte, A. W. and D. Tautz 2010 Understanding the onset of hybrid speciation. Trends in Genetics 26: 54–58.

Otto, S. P. and M. C. Whitlock, 1997 The probability of fixation in populations of changing size. Genetics 146: 723–733.

Palamara, P. F., T. Lencz A. Darvasi and I. Pe’er, 2012 Length distributions of identity by descent reveal fine-scale demographic history. The American Journal of Human Genetics 91: 809–822.

Pan, Q., F. Ali X. Yang J. Li and J. Yan 2012 Exploring the genetic characteristics of two recombinant inbred line populations via high-density snp markers in maize. PLoS ONE 7.

Payseur, B. A. and L. H. Rieseberg, 2016 A genomic perspective on hybridization and speciation. Molecular Ecology.

Pool, J. E. and R. Nielsen 2009 Inference of historical changes in migration rate from the lengths of migrant tracts. Genetics 181: 711–719.

Roesti, M., A. P. Hendry, W. Salzburger and D. Berner 2012 Genome divergence during evolutionary diversification as revealed in replicate lake-stream stickleback population pairs. Molecular Ecology 21: 2852–2862.

Sedghifar, A., Y. Brandvain and P. Ralph 2016 Beyond clines: lineages and haplotype blocks in hybrid zones. Molecular Ecology pp. n/a–n/a.

Singhal, S., E. M. Leffler, K. Sannareddy I. Turner O. Venn A. M. Hooper, A. I. Strand, Q. Li B. Raney C. N. Balakrishnan, S. C. Griffith, G. McVean and M. Przeworski 2015 Stable recombination hotspots in birds. Science 350: 928–932.

Smagulova, F., I. V. Gregoretti, K. Brick P. Khil R. D. Camerini-Otero, and G. V. Petukhova, 2011 Genome-wide analysis reveals novel molecular features of mouse recombination hotspots. Nature 472: 375–378.

Stam, P., 1980 The distribution of the fraction of the genome identical by descent in finite random mating populations. Genetical Research 35: 131–155.

Stemshorn, K. C., F. A. Reed, A. W. Nolte, and D. Tautz 2011 Rapid formation of distinct hybrid lineages after secondary contact of two fish species (cottus sp.). Molecular Ecology 20: 1475–1491.

